# BiFCo: Visualising cohesin assembly/disassembly cycle in living cells

**DOI:** 10.1101/2023.01.21.525018

**Authors:** Emilio González-Martín, Juan Jiménez, Víctor A. Tallada

**Affiliations:** Centro Andaluz de Biología del Desarrollo, Universidad Pablo de Olavide-Consejo Superior de Investigaciones Científicas-Junta de Andalucía, Carretera de Utrera Km1, 41013 Seville, Spain

## Abstract

Cohesin is a ring-shaped protein complex highly conserved in evolution that is composed in all eukaryotes of at least two SMC proteins (Structural Maintenance of Chromosomes) SMC1 and SMC3 in humans (Psm1 and Psm3 in fission yeast), and the kleisin RAD21 (Rad21 in fission yeast). Mutations in its components or its regulators cause genetic syndromes (known as cohesinopathies) and several types of cancer. It has been shown in a number of organisms that only a small fraction of each subunit is assembled into complexes. Therefore, the presence of an excess of soluble components hinders dynamic chromatin loading/unloading studies using fluorescent fusions *in vivo*. Here, we present a system based on bimolecular fluorescent complementation in the fission yeast *Schizosaccharomyces pombe*, named Bi-molecular Fluorescent Cohesin (BiFCo) that selectively excludes signal from individual proteins to allow monitoring the complex assembly/disassembly within a physiological context during a whole cell cycle in living cells. This system may be expanded and diversified in different genetic backgrounds and other eukaryotic models, including human cells.

## Introduction

The cohesin complex has long been attracted a big deal of attention since it has been tightly associated with the structure, function and segregation of eukaryotic genomes. Its interaction with chromatin is critical for establishing topologically associated domains, known as TADs, in order to maintain the three-dimensional structure of genomes for the regulation of gene expression over long distances. Furthermore, it is also necessary for preserving the fidelity of other intrinsic DNA processes such as replication and repair, transcription, condensation and cohesion, which are essential for maintaining genomic stability (Merkenschlager and Nora 2016; Osadska *et al*. 2022). In fact, mutations in some of its components or its regulators cause genetic syndromes (known as cohesinopathies) and several types of cancer in humans (reviewed in (De Koninck and Losada 2016; Di Nardo *et al*. 2022). Cohesin is a ring-shaped complex highly conserved in the evolution from yeast to human. It is composed in all organisms of at least two SMC proteins (Structural Maintenance of Chromosomes) SMC1 and SMC3 (Psm1 and Psm3 in fission yeast), and the kleisin MCD1/SCC1/RAD21 (Rad21 in fission yeast) (Peters *et al*. 2008; Yatskevich *et al*. 2019). When assembled, each of the SMC proteins folds back on itself through a domain known as the “hinge”, bringing its N-terminus and C-terminus together to form a domain known as the “head”. The hinge domains of both proteins interact with each other and the α-kleisin Rad21 links the head domains together to close the ring (Haering *et al*. 2002; Gruber *et al*. 2003) (fig. 1A). The proper functions of cohesin depend to a large extent on the timing of association/dissociation dynamics from chromatin in tight coordination with the progress of the cell cycle. It starts to be recruited to DNA in the G1 phase and progressively accumulates through to the end of S phase, establishing its cohesive action during DNA replication (Guacci *et al*. 1997; Michaelis *et al*. 1997; Uhlmann and Nasmyth 1998; Bernard *et al*. 2008). Subsequently, it is particularly important to remove it timely from mitotic chromosomes in metaphase/anaphase transition to allow physical separation of sister chromatids (Guacci *et al*. 1997; Michaelis *et al*. 1997). The release from DNA in vertebrate cells occurs in two steps. Bulk of chromosomal arms cohesin is removed in prophase (in a series of events called prophase pathway); but centromeric cohesin remains associated until the metaphase to anaphase transition (Losada *et al*. 1998; Waizenegger *et al*. 2000; Losada *et al*. 2002). Centromeric cohesin is then released by a highly conserved mechanism -finely controlled by mitotic regulators-leading to the proteolysis of Rad21 kleisin by a protease known as separase (Cut1sp/ESP1sc/ESPL1hs). Separase is otherwise kept inactive during most of the cell cycle by physical association with another protein called securin (Cut2sp/PDS1sc/PTTG1hs). As soon as the chromosomes in the metaphase plate are properly captured by the mitotic spindle, the Anaphase Promoting Complex (APC/C) ubiquitinates and targets securin for degradation. This allows the activation of separase, which in turn cleaves Rad21 (Funabiki *et al*. 1996; Uhlmann *et al*. 1999; Uhlmann *et al*. 2000; Hauf *et al*. 2001). Rad21 cleavage is essential for a faithful mitosis in many organisms, including yeasts, *Xenopus*, mouse and human. However, it is remarkable that biochemical analyses show that only a small fraction of cohesin’
ss proteins are found to be associated to chromatin before anaphase (Losada *et al*. 1998; Schmiesing *et al*. 1998; Darwiche *et al*. 1999; Tóth *et al*. 1999; Tomonaga *et al*. 2000; Weitzer *et al*. 2003). This is usually not a big concern for most *in vitro* studies as the soluble fraction is washed away in breaking the cells. However, the high soluble amount of each component in intact cells hampers *in vivo* dynamic studies by fluorescent tagging. In this work, we have developed a system in fission yeast, based on bimolecular fluorescent complementation, that allows us to follow only the fraction of these proteins that are physically associated with each other in living cells. Hereinafter, we refer to it as BiFCo (<u>Bi</u>molecular <u>F</u>luorescent <u>Co</u>hesin). Bimolecular fluorescent complementation involves fusing two parts of a fluorescent protein (FP) to two cellular proteins respectively. Each part of the FP is not able to emit fluorescence on its own. However, if the proteins of interest physically interact with each other, the FP halves complement in *trans* and fluorescence emission is restored (Nagai *et al*. 2001; Hu *et al*. 2002). In this study, we have fused the two parts of Venus YFP (referred to as n-YFP and c-YFP respectively) to the C-terminal ends of Psm1 and Rad21 respectively. This system allows us to monitor the *in vivo* dynamics of cohesin loading and unloading at different stages of the cell cycle and different regions of chromosomes.

**Figure1.**
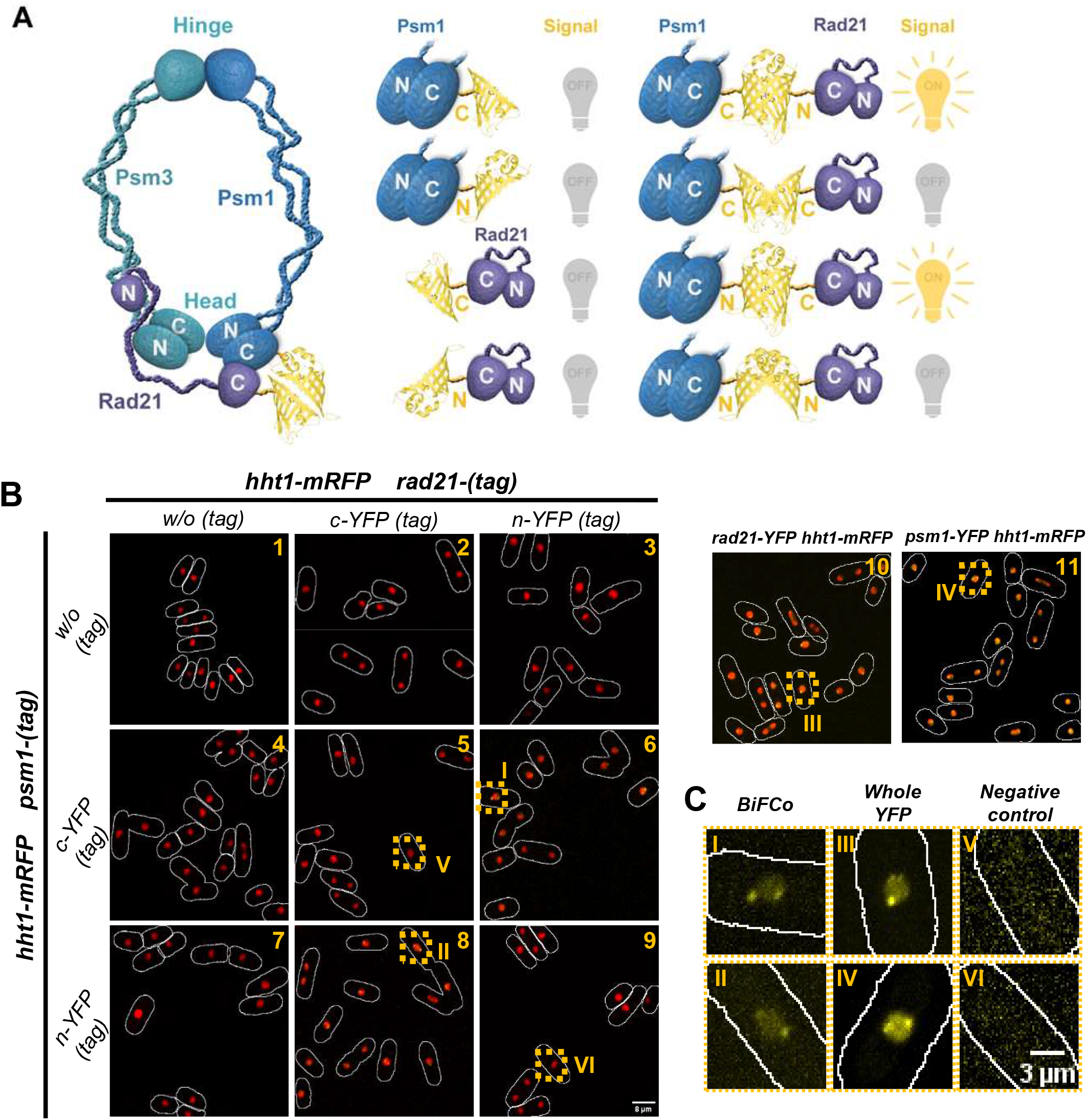
Bi-Molecular Fluorescence Complementation into the cohesin complex. A) Cohesin complex cartoon that shows where YFP moieties are fused (left). In that configuration, only indicated combinations are expected to complement and recover fluorescence capacity (right). B) All generated strains represented in A where crossed into histone Htt1-mRFP strain in order to co-localise BiFCo signal within chromatin. Micrographs presented are sum projections of 21 slices with a Z-step of 0.2 microns. Only whole YFP fusions (upper right panels 10 and 11) and YFP complementing combinations (panels 6 and 8) give detectable yellow signal. Scale bar: 8 μm. C) Representative insets enlargements from section B of YFP channel alone (roman numbers labels). BiFCo signal (I and II) along with positive (III and IV) and negative (V and VI) controls are shown. Scale bar: 3 μm.

## Results

### Bi-molecular Fluorescence complementation at the cohesin head domain

It is very well stablished that the C-terminal ends of Rad21 and Psm1 bind to each other in close proximity of Psm3 to form the cohesin head domain (Haering *et al*. 2002; Haering *et al*. 2004). Thus, we chose these ends for gene tagging at native loci with the bimolecular complementation system as, *a priori*, it is highly likely that the YFP parts will come close enough in space when these proteins interact to form the ring. In addition, the promoter region of these genes remains undisturbed to keep wild type expression regulation of the tagged genes. We tagged C-termini of both Rad21 and Psm1 respectively with the two YFP parts to generate all combinatory by mating and tetrads pulling. These include the reciprocal experimental set-up and the respective negative controls (fig. 1A). We first tested the viability of all strains within the standard growth temperature range of fission yeast wild-type and temperature conditional mutants between 20° and 36°, as compared with the whole YFP control taggings and wt untagged control (supp. fig. 1). All strains grow as the wild-type at most temperatures. Only the fusion of Psm1 with the N-terminal moiety of YFP seems to affect cell viability, albeit only at 36°. Viability though is fully restored when the complementary C-YFP part is present in Rad21 counterpart (experimental configuration) (sup. fig. 1). We then tested the fluorescent signal obtained in live cells both in the two experimental combinations and in the respective negative controls. As expected, no signal is obtained in strains carrying only one part -or two equal parts-of YFP as the untagged control (fig. 1B: panels 1,2,3,4,5,7,9 and fig 1C: panels V, VI). Fluorescent signal was only observed in the two combinations carrying complementing parts of YFP (Rad21-nYFP, Psm1-cYFP and Rad21-cYFP, Psm1-nYFP) (fig. 1B: panels 6,8 and fig. 1C: panels I and II), as well as in the positive controls of proteins carrying a traditional fusion with the whole YFP (fig. 1B: panels 10,11 and fig. 1C: Panels III, IV)). However, while the signal of the complete fusions is very evident in both the nucleoplasm and specific foci, the BiFCo signal is much fainter in the nucleoplasm and concentrated mainly to specific foci. The fluorescence intensity difference suggest that BiFCo is effectively able to lighten exclusively the associated fraction within the complex over total individual components within a living cell.

### BiFCo subnuclear locations *in vivo*

Immunofluorescence, immunoblotting and numerous global *in vitro* genomic studies, using different methods to identify cohesin’
ss immunoprecipitated chromatin (ChIP), have mapped the distribution of cohesin across the genomes. These maps show a high concentration of this complex in telomeric and pericentromeric heterochromatin regions of the chromosomes in yeast and animal cells as well as in specific chromosome arms regions. These very frequently coincide with convergent transcription sites in yeast (Blat and Kleckner 1999; Tanaka *et al*. 1999; Tomonaga *et al*. 2000; Glynn *et al*. 2004; Gullerova and Proudfoot 2008; Koch *et al*. 2008) and regions where CTCF zinc finger protein is located in mammals (Parelho *et al*. 2008), including humans (Holzmann *et al*. 2019). Thus, we tested whether the most prominent fluorescent foci observed by BiFCo correspond to these pools. We co-localised respectively the BiFCo signal *in vivo* with well-established fluorescent markers of telomeres (Taz1 (fig.2 inset I)), centromeres (Mis6 (fig.2 inset II)) and the spindle pole body (Sid4 (fig.2 inset III)), since centromeres remain clustered at the latter structure throughout most of the *S. pombe* cell cycle. In all cells, the brightest BiFCo foci co-localise with either fluorescent marker signal. This confirms that it is possible to follow dynamically the accumulation of cohesin associated with centromeres and telomeres at the same time (fig. 2) and it could indeed serve in interphase as a simultaneous marker of these chromosomal regions in living cells. Apart from these sites, the system is sensitive enough to detect numerous smaller foci within the nuclei that likely correspond to accumulation loci along the chromosome arms (fig.2 inset IV). Thus, BiFCo pattern *in vivo* results strikingly similar to that found by chromosome spreads immunofluorescence against Rad21 in fixed cells where soluble cohesin components are washed away and it only remains visible the chromatin-bound cohesin fraction (Ciosk *et al*. 2000; Schmidt *et al*. 2009).

**Figure 2.**
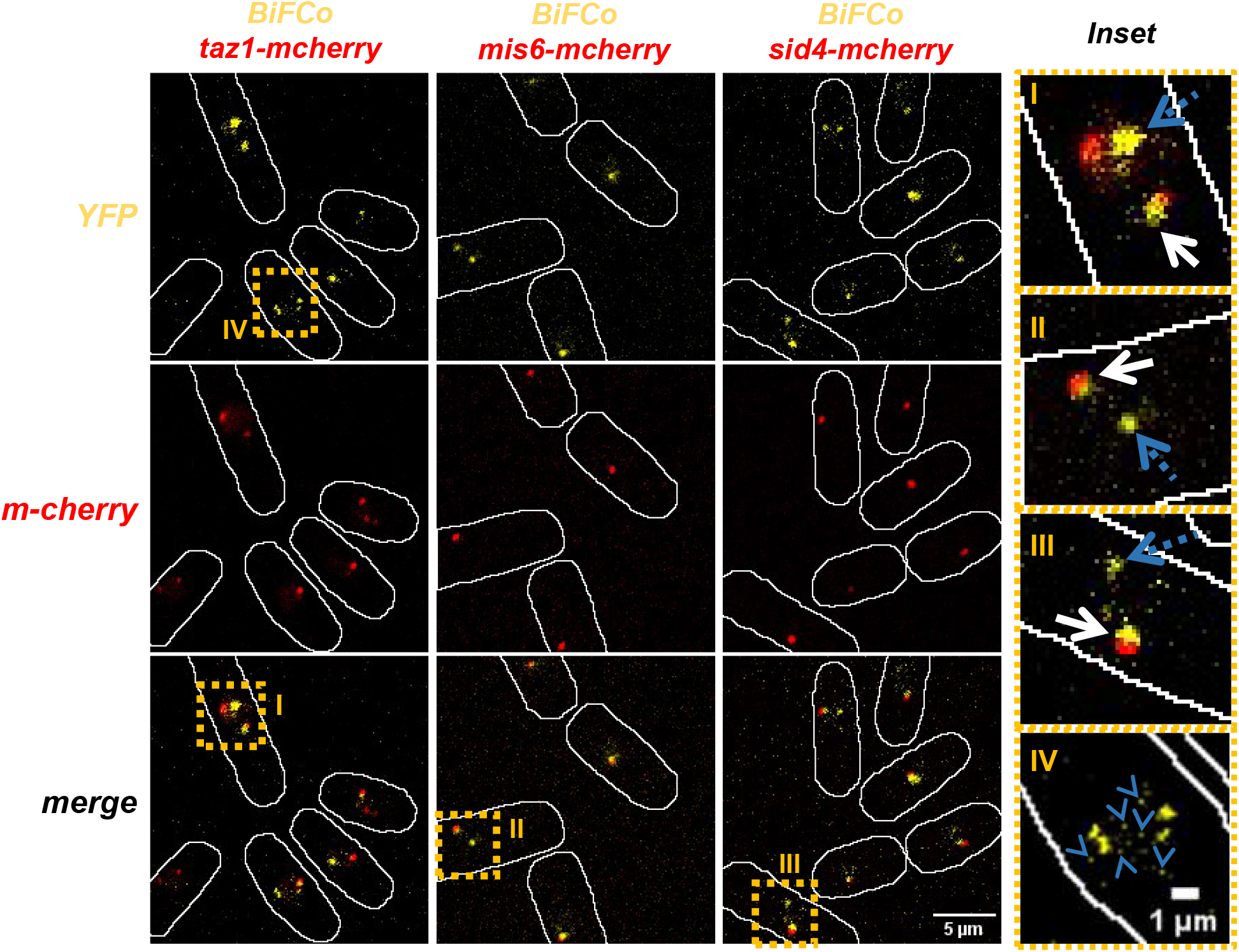
Co-localization of BiFCo foci. We crossed our Bi-molecular fluorescent cohesin system into Taz1-mcherry, Mis6-mCherry and Sid4-mCherry backgrounds to co-localise telomeres, kinetochores (centromeric region) and the SPB respectively. High contrast images of maximal projections of 21 slices with a Z-step of 0.2 microns are presented. Single channel and merged images are shown. Scale bar: 5 μm. Inset enlargements of merged images are shown on the right (I, II and III roman numbers). In all cases some BiFCo main signal foci co-localise with either of the red markers (white continuous arrows) but others do not (blue dotted arrows). Bigger enlargement of high contrast merged images (IV) allows to distinguish also numerous secondary foci (blue arrowheads). Scale bar: 1 μm. Taken together this data indicate that the more intense BiFCo signal correspond to cohesin accumulation at centromeres and telomeres and secondary foci represent assembled cohesin at chromosomal arms loci.

### Dynamic analysis of assembled cohesin

The function of cohesin complex is essential for reversible cohesion and the correct segregation of sister chromatids but it is also fundamental for DNA replication, the regulation of gene expression and the definition of topologically associated domains (TADs) and consequently for the maintenance of genomic stability (Nasmyth and Haering 2005; Losada 2008; Peters *et al*. 2008). This highly conserved dynamics in eukaryotes is achieved by regulating the loading and unloading of cohesin on chromatin. The release of sister chromatids to be separated to opposite poles in mitosis is achieved by the degradation of Rad21 kleisin, by the action of the Cut1 protease. We therefore asked whether this proteolysis of Rad21 can be monitored by the decay of the BiFCo signal in time-lapse experiments in living cells (fig.3A). To graphically represent this dynamics, we quantified the nuclear fluorescence at every time lapse in dividing nuclei and normalised by the fluorescence of the time lapse in which we observed the characteristic oblong shape of the nucleus at the beginning of anaphase B (labelled as time 0 in the figures). As shown, there is an abrupt drop of the signal at an interval of 6 min before and 20 min after the normalisation point and a simultaneous recovery of fluorescence in the two daughter nuclei corresponding to *de novo* loading in G1 (fig. 3C). These data show that the proteolytic break down and re-association of Rad21-Psm1 interaction can be timely followed in living cells. However, even at the lowest level of fluorescence, a faint nuclear signal is still detected above the background noise. This signal might correspond to assembled free or non-cohesive rings (Huis in ‘t Veld *et al*. 2014) or single-chromatid loaded rings whose Rad21-Psm1 interaction may not be dislodged in mitosis (Huang *et al*. 2005; Schmidt *et al*. 2009; Nasmyth 2011; Eng *et al*. 2015). Next, we tested the robustness of the BiFCo system over time by following two mitotic divisions (fig. 3B). In order to gather a significant number of nuclei going through two mitoses in the same field of view and to minimise signal variability by bleaching effects and photo-toxicity, we synchronised cells by using the conditional *cdc25*.*22* mutant background (Russell and Nurse 1986). This mutant arrests the cell in late G2 at high temperature (36°), but it enters normally into mitosis when released at the permissive temperature of 25° under the microscope. As shown in figure 3D and supplementary video 1, BiFCo signal can be followed for at least two Rad21 cleavage cycles, indicating that cohesin dynamics can be monitored for a very long period (8 hours) and more than one assembly-disassembly cycle.

**Figure 3.**
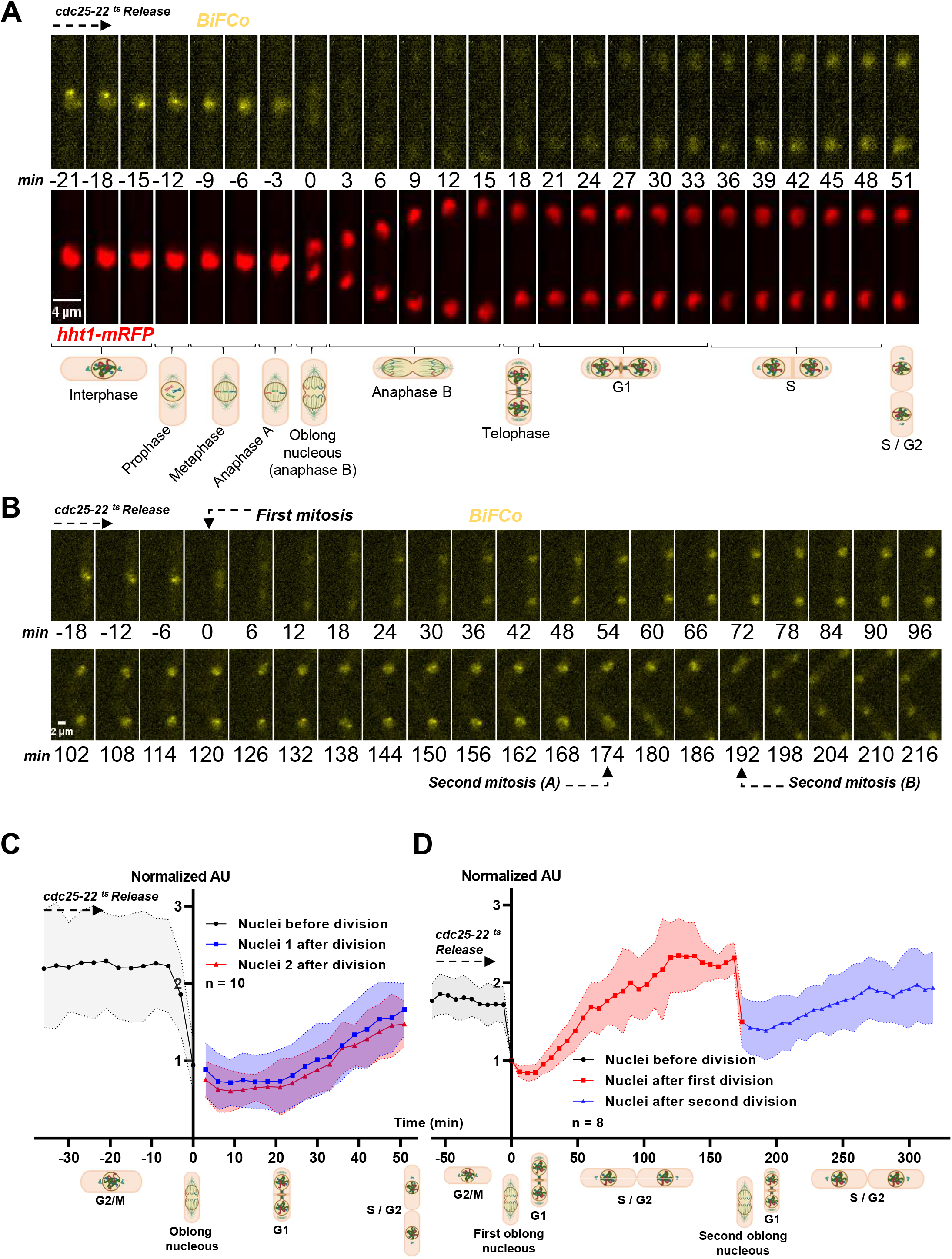
Cohesin’ss cycle dynamics. A) Individual cells bearing the BiFCo system into Htt1-mRFP background (as those in figure 1B panel 8) were followed in 3 minutes time-lapse experiments in the transition from G2 to G1/S phase to assess predicted degradation in mitosis. Sum Projections frames are shown above the schematic representation of cell cycle phases. Scale bar: 4 μm. B) Synchronized cdc25-22 BiFCo cells were arrested for 4 hours at restrictive temperature and were released and monitored under the microscope for eight hours at permissive temperature to assess signal recovery and decay over two nuclear divisions. Scale bar: 2 μm. C) Fluorescent signal quantification over time. Ten individual nuclei divisions from section A were analysed and average fluorescence at each time lapse is represented. In order to compare the cell cycle stage for every nuclei as close as possible, we set out first time lapse of oblong nucleus shape (beginning of anaphase B) as time 0. We used the average level of fluorescence of oblong nuclei minus average background level at the same time point to normalise fluorescence for each time lapse (fNuclei-fBkd / fOblong each time lapse-fBkd each time lapse). Thus, normalised fluorescence of 1 represents the average intensity at the beginning of anaphase B. D) Signal quantification (as in C) from section B over two consecutive mitotic events.

### Total vs assembled cohesin components

As explained above, it has been traditionally difficult to follow the assembly and disassembly dynamics of cohesin rings by fluorescence throughout the cell cycle or in experimental situations such as cohesin regulators’ mutant backgrounds. This is because there is only a proportion of its components forming the complex, relative to the total amount of each component separately. Out of those, not all are bound to chromatin; but even out of the bound ones, not all are degraded in mitosis (Losada *et al*. 1998; Schmiesing *et al*. 1998; Darwiche *et al*. 1999; Tóth *et al*. 1999; Tomonaga *et al*. 2000; Weitzer *et al*. 2003; Schmidt *et al*. 2009; Holzmann *et al*. 2019). Actually, separase cleaves only 10% of cohesin complexes in mammalian cells (Waizenegger *et al*. 2000; Koch *et al*. 2008) and *Drosophila* cells can tolerate an artificial decrease of 80% of total Rad21 levels before exhibiting cohesion defects (Carvalhal *et al*. 2018). The results above suggest that most –if not all-of the Rad21 cleaved population correspond to the fraction bound to Psm1. Thus, we compared the fluorescence of the BiFCo system with traditional entire YFP fusions in the same time-lapse experiment with identical settings (fig. 4A, supp. fig 2 and supp. Video 2). Since we had assessed before that the two daughter nuclei behave in the same way, in terms of fluorescence recovery, we monitored in this case only one nucleus from each division to simplify the plotting. At the G2-M transition, fluorescence levels of the total population of Psm1 are 3,8 fold higher than Rad21 and both signals become 3,2 and 12 fold higher respectively than that observed for the BiFCo system. This suggests that only about 30% of Rad21 and 8% of Psm1 are interacting close enough to each other into the BiFCo complex. Both proteins undergo a drop in fluorescence in mitosis, as expected when the content of one nucleus is split into two. Rad21 average signal drops by 50,1% from the highest level in G2 to the lowest level in late anaphase/early G1. Studies in fission yeast’
ss cell lysates immunoblots have reported that less than 5% of Rad21 is cleaved in mitosis (Tomonaga *et al*. 2000). So the further decay from the 50% expected may be very subtle to be detected as an average fluorescence of a number of cells. Alternatively, it might be compensated with new protein import into the nucleus. Actually, Psm1 fluorescence drops only by 29,2%; clearly less than expected (50%), suggesting a rapid import of this protein into the nucleus during anaphase which might compensate fluorescence loss at nuclear splitting. In contrast to the whole fusions, the average BiFCo signal decreases by 68% in this period, making this unloading transition very evident visually and quantifiable (fig. 4B). This drop likely accounts for cohesin disassembly, both dependent and independent of Rad21 proteolysis, although the latter seems to be a small fraction by immunoblotting and chromosome spread analyses (Schmidt *et al*. 2009). Thus, these data show that mitotic detachment of cohesin can be reliably monitored in living cells.

**Figure 4.**
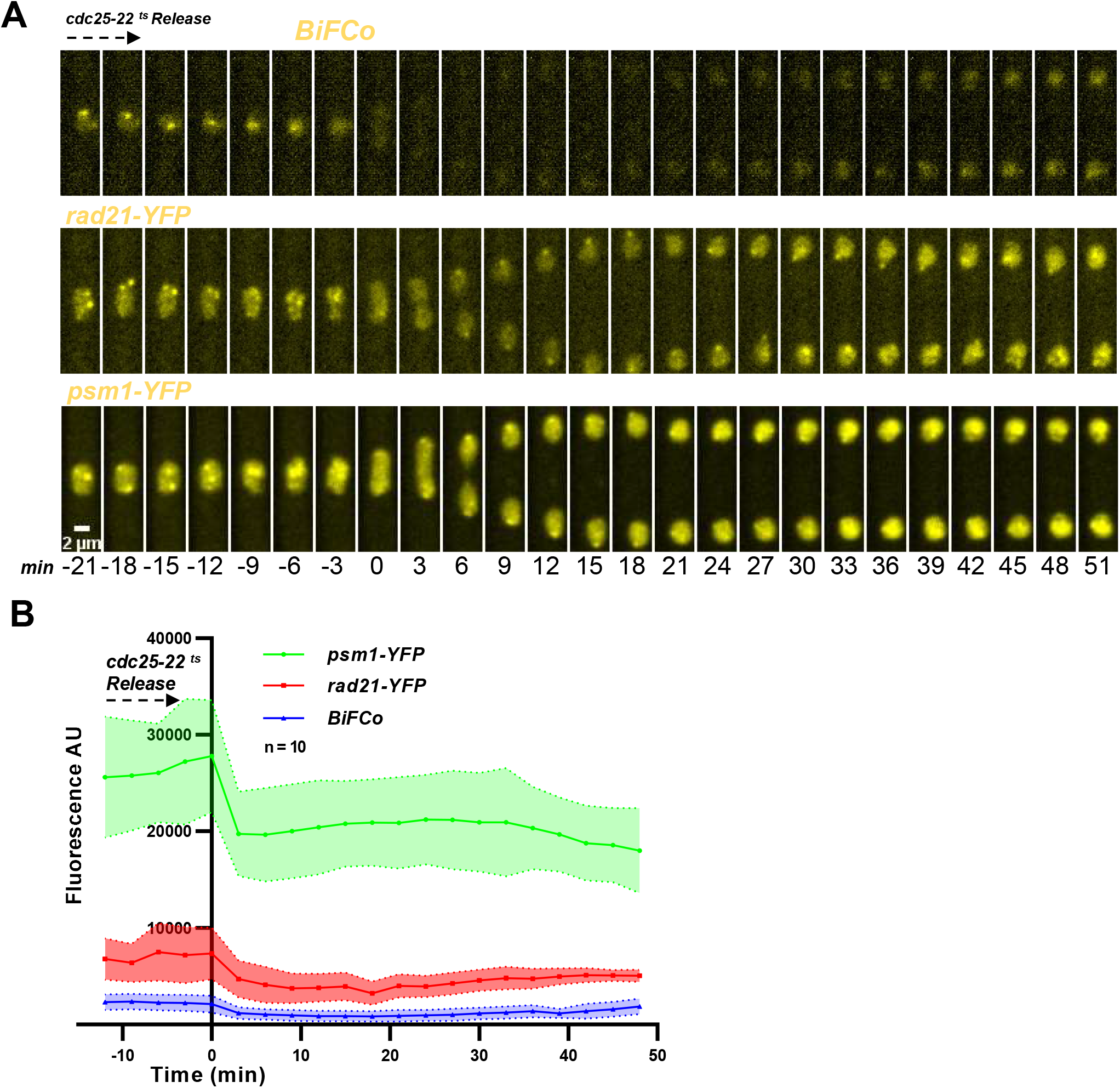
Total vs assembled cohesin components. A) Whole YFP fusions to Rad21 and Psm1 respectively, together with the BiFCo strain, where mounted onto the same coverslip but spatially separated. This is achieved by sticking the cells to separate and confined soybean lectin drops. In order to compare fluorescent signal in cells getting into mitosis, time-lapse microscopy (travelling over the three strains in each time point) was performed under identical illumination/capturing set up. Scale bar= 2 μm. Note that brightness and contrast levels of each strain presented in this figure are adjusted separately in order to distinguish nuclear foci in all strains. Real signal comparison with identical brightness and contrast levels can be seen in section B, supplementary figure 2 and supplementary video 1. B) Average fluorescence intensity and standard deviation (arbitrary units) are plotted. Ten nuclei for each strain were analysed in the same cell cycle stage. The equivalent background area at each time point for each cell was subtracted from the actual signal within the nucleus. It should be noted that the BiFCo system plotting appears to be flat due to the large increase in the scale of the arbitrary fluorescence units when individual whole fusion proteins are plotted along.

### Qualitative and Quantitative BiFCo responses to cell cycle signalling

In order to confirm that BiFCo signal is proportional to the amount of assembled complex and it responds to cell cycle cues for cohesin loading and unloading dynamics, we impaired known regulators of these transitions and blocked the cell cycle in different stages. As mentioned above, the cohesin release in mitosis is mostly dependent on Rad21 proteolysis, which breaks up Psm1-Rad21 interaction. Thus, we monitored BiFCo in a mutant background in which Rad21 cannot be degraded. In a wild-type situation, entrance into mitosis activates the APC/C that targets Cut2 for destruction. This activates Cut1 protease, that in turn, cleaves Rad21 (Cohen-Fix *et al*. 1996; Funabiki *et al*. 1996). Thus, we expressed the BiFCo system into *cut9-665* background. This temperature-sensitive allele, at the restrictive temperature of 36°C, renders the APC/C inactive and therefore Rad21 cannot be degraded downstream (Samejima and Yanagida 1994). We switched this strain to restrictive temperature for four hours. Then we mounted the cells for live microscopy -keeping restrictive temperature under the microscope-to take time-lapse images every three minutes for two more hours. As it can be seen in figure 5A, cells that end up with segregation defects after septation, do not show the drop in BiFCo signal when going through mitosis. Complementarily, we ensured that the majority of the observed BiFCo signal is dependent on the loading of the complex on chromatin in interphase. For this purpose, we observed the complex together with a red chromatin marker (Htt1-mRFP) in the temperature-sensitive *mis4-242* mutant background. At the restrictive temperature of 36°, this mutant is defective in the loading function and unable to maintain chromosome association (Tomonaga *et al*. 2000) (Bernard *et al*. 2008). We observe a fluorescent signal similar to the wild-type background at the permissive temperature (25°). Even the foci of centromeres and telomeres are visible (fig. 5B upper panels). However, when Mis4 function is impaired at 36°, cohesion defects are evident as judged by the chromatin marker (red channel) and the BiFCo signal is extremely reduced to hardly detectable levels (fig. 5B lower right panel). Apart from the implications of this result in the BiFCo signal dependence on Mis4 function, the fact that the fluorescent signal almost disappears also implies that the interaction in *trans* of both parts of YFP does not abnormally stabilise the complex, but is reversible and responds to the natural process of cohesin disassembly (see discussion). These data together show that the BiFCo signal is dependent on Mis4 loading function and its severe drop in mitosis is a direct consequence of Rad21 proteolysis induced by the APC/C. Complementarily, we asked whether the fluorescent bipartite system is able to detect quantitative phenotypes in cohesin ring assembly. In fission yeast, cohesin recruitment to chromatin begins in G1. In this phase, the loading/unloading dynamics reaches a stationary state, as in human cells (Guacci *et al*. 1997; Michaelis *et al*. 1997). Further accumulation and stabilization occurs in S-phase, which lasts through to G2 phase until mitosis (Uhlmann and Nasmyth 1998; Bernard *et al*. 2008). Therefore, we arrested the cell cycle at different stages and quantified the BiFCo signal in 50 random nuclei for each stage. We assessed an asynchronous wild type control, nitrogen deprived cells (G1 arrest), *cdc10-129* mutant (START arrest), Hydroxyurea (S phase arrest) and *cdc25*.*22* mutant (G2/M arrest). As expected, the asynchronous population - in which only about 10% of cells are in mitosis - showed no significant difference in the average signal intensity with G2-arrested-cells. However, we detected significant increasing signal from asynchronous cells compared respectively to a G1 arrest at 4 hours of nitrogen starvation (1,7 fold), G1/S transition arrest (2,8 fold), and to an early S-phase block by hydroxyurea (4,1 fold) (fig. 5C, D). These data show that BiFCo can be used as a quantitative marker of chromatin loaded cohesin but it also have interesting implications to study cohesin assembly and its coordination to the cell cycle.

**Figure 5.**
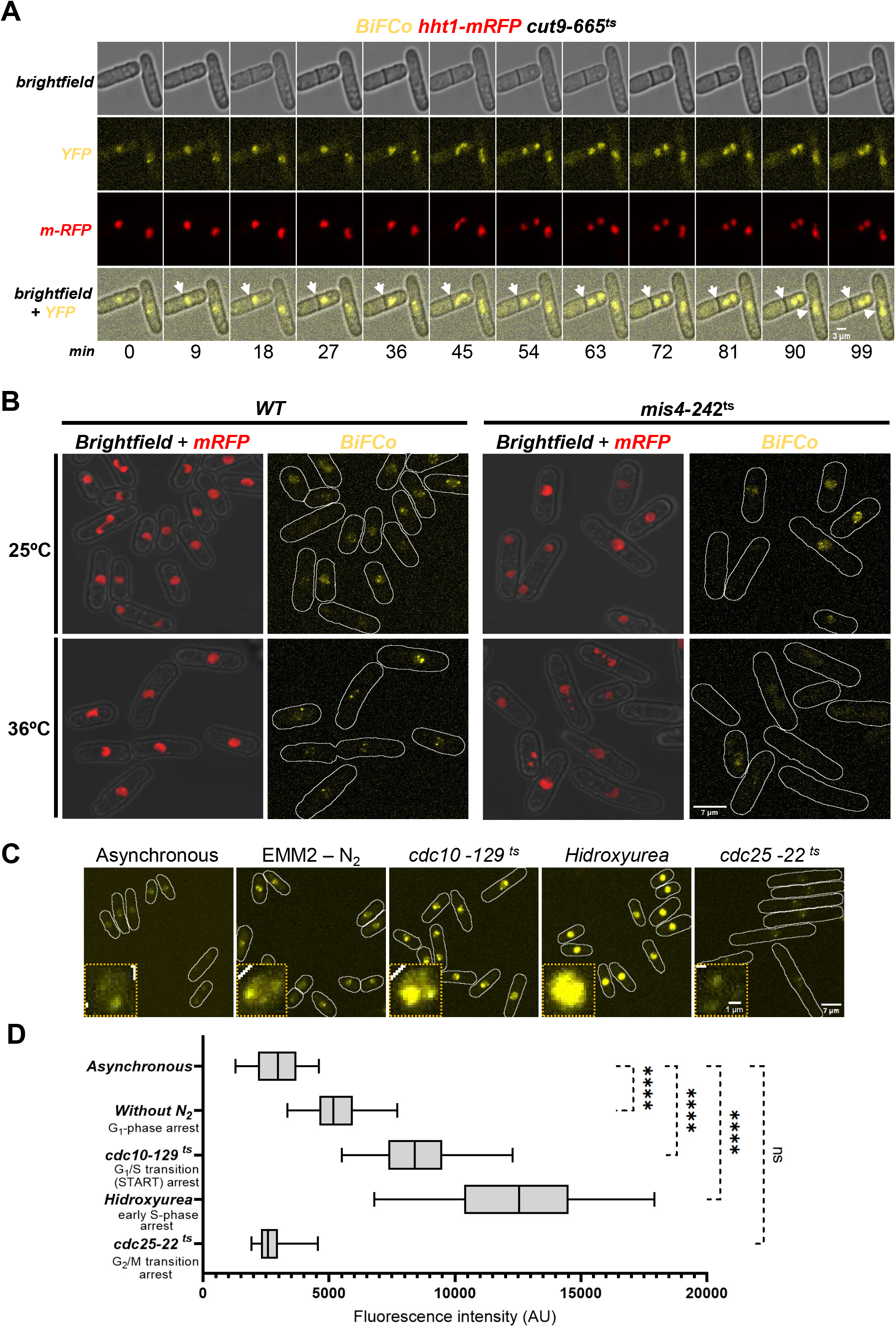
BiFCo signal depends on Mis4 loader function and the mitotic decay depends on Rad21 degradation. A) The BiFCo tagging was expressed into the temperature-sensitive cut9-665 mutant background. At high temperature, the APC/C function is abolished, so Rad21 is not cleaved. We cultured these cells until early log phase at permissive temperature of 25° and shifted them to 36° for 4 hours before mounting them for time lapse microscopy, keeping the sample at high temperature under the microscope. Contrary to wild type mitosis, YFP signal from the assembled complex do not decays in cells that go through anaphase. Time lapse interval is indicated at the bottom: 9 min (Scale bar= 3 μm). B) BiFCo complex, together with Htt1-mRFP (red chromatin marker), was expressed into a mis4+ and mis4-242ts backgrounds. Exponentially growing cultures at 25° and shifted cultures to 36° for four hours were processed for live cell microscopy. Maximal projections from twenty-one Z-stacks every 0,2 μm are shown for both strains and temperatures under identical settings. Scale bar: 7 μm. Fluorescence from the assembled complex is fully dependant on Mis4 function. C) BiFCo maximal projections (21 slices) from either asynchronous cells, nitrogen starved cells for 4 hours, cdc10-129 arrested cells at 36º for four hours, hydroxyurea treated cells (10 mM, 4 hours) or cdc25-22 arrested cells at 36º for four hours. Scale bar: 7 μm. The inset shows a representative nucleus enlargement in each case. Scale bar: 1 μm. Images are taken and adjusted identically for brightness and contrast D) Box and violin distributions for signal quantification of 50 random nuclei in each condition. We used ordinary one-way ANOVA comparison against asynchronous control distribution to test statistical significance (GraphPad Prism 9.0.2 software. Dunnett’s multiple comparisons test). Confidence interval p<0,05. ns: non-significant. Asterisks indicate p-value <0,0001

## Discussion

The cohesin complex has attracted much attention over the last three decades, during which we have learned about the essential and evolutionarily highly conserved functions that performs in all the architecture, expression and segregation of eukaryotic genomes. Defects in the assembly of complex’
ss subunits and in the dynamics of loading and unloading onto the chromosomes are a common cause of genetic diseases in humans (Remeseiro *et al*. 2013). Therefore, the study of these processes helps us to understand and seek treatment for these diseases. Very important structural features and dynamics of the cohesin complex have already been discovered in living cells by fluorescence techniques such as FRAP and FRET (Mc Intyre *et al*. 2007; Holzmann *et al*. 2019). Although technical limitations in sensitivity and temporal resolution make undetectable possible transient cohesin structural changes and long-term loading and unloading transitions. In addition, the use of complete fluorescent proteins fusions make indistinguishable the signal of total components from the assembled fraction. In fact, the vast majority of cohesin loading, structure and regulation studies have been carried out *in vitro*, in reconstituted reactions, analysing cell lysates or fixed chromosome spreads immunofluorescence (Bernard *et al*. 2008; Xiang and Koshland 2021). In this work, we have developed the proof of principle of bimolecular fluorescent cohesin (referred to as BiFCo), which selectively discriminates the assembled complex within a physiological context; excluding the soluble fraction of individual proteins present in living cells. We show that separate fusions of the two complementary parts of vYPF to Psm1 and Rad21 respectively, only give rise to fluorescence when these proteins interact with each other as part of the complex. Consistent with *in vitro* cohesin-associated chromatin immunoprecipitation analyses (ChIP-seq, ChIP-chip), fixed chromosome spreads and FRET studies (Blat and Kleckner 1999; Tanaka *et al*. 1999; Mc Intyre *et al*. 2007; Schmidt *et al*. 2009), we detect the BiFCo signal at major sub-nuclear foci in interphase, corresponding to centromeric and telomeric regions of the chromosomes. Furthermore, the sensitivity of this system also allows detection at minor foci that likely correspond to clusters along the chromosome arms.

The use of the bimolecular complementation approach is particularly useful for the dynamic study of protein interactions *in vivo* in the same cell over time. However, in some cases, it faces two potential limitations. On the one hand, the folding time of the two parts of YFP fused to their respective baits and the time needed for the association of fluorescent fragments and chemical reactions leading to the fluorophore production, may be limiting when the proteins turnover in the given biological process to address is very fast. On the other hand, it has been described that the re-composition of the two parts of YFP *in trans* becomes very stable once is produced, making it difficult to study *in vivo* protein interactions that must be naturally reversible (reviewed in (Kerppola 2009)). However, in the experimental system presented here, these limitations are inherently overcome by the biology of cohesin itself. The fact that there exists a large pool of free components present in the nucleus, in addition to pre-assembled complexes, suggests that the cell is provided with a natural reservoir of complexes and soluble partners that are ready to be loaded or coupled *de novo* to ensure a rapid reloading in G1 phase. Thus, BiFC-tagged proteins should have sufficient time for synthesis and folding before they are assembled in the next cycle. This is illustrated in our analysis by the rapid onset of fluorescence recovery from the lowest intensity time lapse; just in the next 20 minutes, coincident with wild type G1 phase. On the other hand, it has been shown in *S. cerevisiae* that once separase cleaves SCC1/Rad21, the resulting C-terminal fragment is rapidly degraded by the N-end rule pathway and it otherwise causes lethality when it is artificially stabilized (Rao *et al*. 2001). The cleavage recognition sites in SCC1/Rad21 are very similar between budding and fission yeast and the amino acids left at the N-terminus of the resulting fragments are also compatible with the degradation system by the N-end rule pathway (Varshavsky 1996; Tomonaga *et al*. 2000). Provided that we do not observe any sign of anaphase progression defects in the BiFCo strain, potentially caused by altered half-life of the Rad21 fragmented products; it is reasonable to consider that this degron system also operates in *S. pombe* and thus the YFP moiety fused to Rad21 C-terminal fragment is quickly degraded after cleavage. So even if the interaction of the two parts of YFP was very stable, in this case it should not pose a problem; as we are in fact monitoring a stable interaction *per se*, which occurs only once between two partners and it is naturally destroyed to reset the system at each cell cycle. In addition, the signal decay to almost undetectable levels in *mis4* defective background suggest that the YFP moieties interaction can also be physiologically broken independently of separase action. Finally, another interesting observation following two complete mitotic cycles of this set-up is that after a long-term exposure (8 hours) of cells to the excitation laser, the G2 intensity levels and fluorescence recovery rate in the second cycle are very similar to the first one. This suggests that the separate parts of YFP do not appear to undergo a significant photobleaching. This sets an advantage to follow the complex over several cell cycles or in model organisms with prolonged cell cycles.

Using full YFP tagging, compared average nuclear fluorescence suggests that total levels of Psm1 in G2 phase are 3,8 times higher than Rad21 levels. This roughly 4:1 stoichiometry actually represents a considerable difference with that observed in budding yeast, where equal protein levels are detected (Mc Intyre *et al*. 2007). In any case, loading and unloading tracking of the complete fusions throughout the cell cycle is not very apparent in either yeast, because of the large background of soluble protein present within the nucleus. In contrast, the absolute BiFCo signal in G2 is 3,2 and 12 times lower than the Rad21 and Psm1 whole YFP fusions respectively. Importantly, BiFCo complex behaves physiologically as expected as the assembly depends on Mis4 loader function and the signal abrupt decay by almost 70% in mitosis makes clearly detectable the cohesion dissolution in living cells. We show that this signal drop is caused by the separase-mediated cleavage of Rad21 kleisin, since this degradation is not observed when APC/C function is inhibited in the *cut9*.*665ts* mutant background. Nevertheless, detectable basal signal levels remain during wild-type anaphase. It might correspond to chromatin independent, single chromatid assembled complex and/or non-cohesive cohesin. In terms of closing the cycle, it is well stablished in yeast that cohesin is recruited on chromatin and stabilised over a period of the cell cycle comprising G1 and S phases. Cell cycle arrest at different points within these phases shows that BiFCo is able to reveal quantitative cohesin-assembly phenotypes in a dynamic range of at least 4-fold over wild-type levels in G2. A short block in G1 by removing the nitrogen source for only 4 hours results in a 1.7-fold increase over G2 signal levels. Interestingly, the assembly/stabilisation process appears to be continuous and cumulative since it keeps increasing up to 2.8-fold if the block occurs slightly later in the G1/S transition (known as START in yeast and “restriction point” in mammals) and up to 4-fold if the cell is arrested early in S-phase by the addition of HU. For the interpretation in the case of HU-mediated arrest, it has to be taken into consideration that HU also induces DNA damage and activates the DNA replication checkpoint. In fact, some cohesin mutants are hypersensitive to several DNA-damaging agents, including hydroxyurea and this drug induces SCC1/Rad21 phosphorylation (Kim *et al*. 2002; SchÄr *et al*. 2004). Therefore, in addition to the loading process that may be maintained while the cell transits through S phase regardless the actual progression is blocked, activation of the replication checkpoint may also contribute to the increased signal that we observe.

Up to our knowledge, this is the first proof of principle set up to visualise cohesin to track assembly/disassembly dynamics within a living cell over a whole cell cycle. We think this system may be expanded and diversified by fusing the complementary parts of suitable fluorescent proteins to different domains and/or to different proteins forming complex and their regulators. The integration of BiFCo with diverse mutant genetic backgrounds may also help in the design of high-content screens and experiments to answer important questions within a living cell context about the intricate functions, dynamics, structural biology and recent loop extrusion models of cohesin (Reviewed in (Higashi and Uhlmann 2022)). This approach could also be tested in other structurally similar SMC complexes such as Condensin and SMC5/6 (Anderson *et al*. 2002; Nasmyth and Haering 2005; Yatskevich *et al*. 2019; Sole-Soler and Torres-Rosell 2020; Kim 2021). Moreover, the high evolutionary conservation of the structure and regulation of these complexes in eukaryotic organisms, together with the advent of CRISPR-Cas editing systems, beckons the possibility of being easily implemented in other biological models from yeast through to human cultured cells.

## Materials and methods

### Media and growth conditions

Standard fission yeast growth media and molecular biology approaches were used throughout as in (Moreno *et al*. 1991). Sporulation agar (SPA) was used for mating and sporulation. Tetrad pulling for segregation analyses was performed in a Singer MSM 400 automated dissection microscope (Singer Instruments). For spot tests, cells were grown to mid-log phase in EMM2 media, cell number/ml was scored in Neubauer’
ss chamber and matched dilutions were calculated for all cultures. Serial five-fold dilutions were plated onto solid media.

### Gene tagging

Gene tagging was performed as described in (Bahler *et al*. 1998) based on pF6aMX6 plasmid series. Candidate tagged strains after transformation were checked by PCR for integration at the expected loci.

### Fluorescence microscopy

Fluorescence images were taken from cells in exponential growth phase in all cases in a Zeiss Axio Observer 7 inverted microscope with Zeiss Plan-Apochromat 63x/1,40 Oil DIC and Alpha Plan-Apochromat 100x/1,46 Oil DIC lenses, coupled to Spinning Disk Confocal Yokogawa CSU-W1 head with excitation lasers and filters from 3i (Intelligent Imaging Innovations). SlideBook 6 software was used for device control and image capturing. Cells were mounted in Biopthec FCS2 chamber for temperature control under the microscope. When fluorescent signal comparison between two or more strains was needed, all strains were stuck next to each other but physically separated onto soybean lectin delimited drops on the coverslip. All images were processed using the open Image J software (Schneider *et al*. 2012). Presented images correspond to maximal or sum projections as indicated in each figure legend.

## Acknowledgments

We thank Dr. Iain Hagan for BiFC tagging plasmids and tagged strains, Dr. Mitsuhiro Yanagida, Dr. Paul Nurse and Dr. Julia Cooper for mutant and tagged strains. Katherina García for assistance in advanced microscopy facility, Modesto Berraquero and Sergio Villa for technical advice and Victor Carranco for excellent lab technical support and digital design advice. Dr. Ignacio Flor-Parra for comments on the manuscript. This work has been funded by Programa operativo FEDER/Junta de Andalucía 2014-2020. Objetivo específico 1.2.3. Grant UPO-1381219 to VAT and Ministerio de Ciencia e Innovación Grant PID2019-111124GB-I00 to JJ. EGM is funded by Pablo Olavide University grant “Ayudas para Investigación Tutorizadas” modalidad B-2 (Rf.: PPI2104).

## References

Anderson, D. E., A. Losada, H. P. Erickson and T. Hirano, 2002 Condensin and cohesin display different arm conformations with characteristic hinge angles. J Cell Biol 156: 419–424.

Bahler, J., J. Q. Wu, M. S. Longtine, N. G. Shah, A. McKenzie, 3rd et al., 1998 Heterologous modules for efficient and versatile PCR-based gene targeting in Schizosaccharomyces pombe. Yeast 14: 943–951.

Bernard, P., C. K. Schmidt, S. Vaur, S. Dheur, J. Drogat et al., 2008 Cell-cycle regulation of cohesin stability along fission yeast chromosomes. Embo j 27: 111–121.

Blat, Y., and N. Kleckner, 1999 Cohesins bind to preferential sites along yeast chromosome III, with differential regulation along arms versus the centric region. Cell 98: 249–259.

Carvalhal, S., A. Tavares, M. B. Santos, M. Mirkovic and R. A. Oliveira, 2018 A quantitative analysis of cohesin decay in mitotic fidelity. J Cell Biol 217: 3343–3353.

Ciosk, R., M. Shirayama, A. Shevchenko, T. Tanaka, A. Toth et al., 2000 Cohesin’s binding to chromosomes depends on a separate complex consisting of Scc2 and Scc4 proteins. Mol Cell 5: 243–254.

Cohen-Fix, O., J. M. Peters, M. W. Kirschner and D. Koshland, 1996 Anaphase initiation in Saccharomyces cerevisiae is controlled by the APC-dependent degradation of the anaphase inhibitor Pds1p. Genes Dev 10: 3081–3093.

Darwiche, N., L. A. Freeman and A. Strunnikov, 1999 Characterization of the components of the putative mammalian sister chromatid cohesion complex. Gene 233: 39–47.

De Koninck, M., and A. Losada, 2016 Cohesin Mutations in Cancer. Cold Spring Harb Perspect Med 6.

Di Nardo, M., M. M. Pallotta and A. Musio, 2022 The multifaceted roles of cohesin in cancer. J Exp Clin Cancer Res 41: 96.

Eng, T., V. Guacci and D. Koshland, 2015 Interallelic complementation provides functional evidence for cohesin-cohesin interactions on DNA. Mol Biol Cell 26: 4224–4235.

Funabiki, H., H. Yamano, K. Kumada, K. Nagao, T. Hunt et al., 1996 Cut2 proteolysis required for sister-chromatid seperation in fission yeast. Nature 381: 438–441.

Glynn, E. F., P. C. Megee, H. G. Yu, C. Mistrot, E. Unal et al., 2004 Genome-wide mapping of the cohesin complex in the yeast Saccharomyces cerevisiae. PLoS Biol 2: E259.

Gruber, S., C. H. Haering and K. Nasmyth, 2003 Chromosomal cohesin forms a ring. Cell 112: 765–777.

Guacci, V., D. Koshland and A. Strunnikov, 1997 A direct link between sister chromatid cohesion and chromosome condensation revealed through the analysis of MCD1 in S. cerevisiae. Cell 91: 47–57.

Gullerova, M., and N. J. Proudfoot, 2008 Cohesin complex promotes transcriptional termination between convergent genes in S. pombe. Cell 132: 983–995.

Haering, C. H., J. Löwe, A. Hochwagen and K. Nasmyth, 2002 Molecular architecture of SMC proteins and the yeast cohesin complex. Mol Cell 9: 773–788.

Haering, C. H., D. Schoffnegger, T. Nishino, W. Helmhart, K. Nasmyth et al., 2004 Structure and stability of cohesin’s Smc1-kleisin interaction. Mol Cell 15: 951–964.

Hauf, S., I. C. Waizenegger and J. M. Peters, 2001 Cohesin cleavage by separase required for anaphase and cytokinesis in human cells. Science 293: 1320–1323.

Higashi, T. L., and F. Uhlmann, 2022 SMC complexes: Lifting the lid on loop extrusion. Curr Opin Cell Biol 74: 13–22.

Holzmann, J., A. Z. Politi, K. Nagasaka, M. Hantsche-Grininger, N. Walther et al., 2019 Absolute quantification of cohesin, CTCF and their regulators in human cells. Elife 8.

Hu, C. D., Y. Chinenov and T. K. Kerppola, 2002 Visualization of interactions among bZIP and Rel family proteins in living cells using bimolecular fluorescence complementation. Mol Cell 9: 789–798.

Huang, C. E., M. Milutinovich and D. Koshland, 2005 Rings, bracelet or snaps: fashionable alternatives for Smc complexes. Philos Trans R Soc Lond B Biol Sci 360: 537–542.

Huis in ‘t Veld, P. J., F. Herzog, R. Ladurner, I. F. Davidson, S. Piric et al., 2014 Characterization of a DNA exit gate in the human cohesin ring. Science 346: 968–972.

Kerppola, T. K., 2009 Visualization of molecular interactions using bimolecular fluorescence complementation analysis: characteristics of protein fragment complementation. Chem Soc Rev 38: 2876–2886.

Kim, K. D., 2021 Potential roles of condensin in genome organization and beyond in fission yeast. J Microbiol 59: 449–459.

Kim, S. T., B. Xu and M. B. Kastan, 2002 Involvement of the cohesin protein, Smc1, in Atm-dependent and independent responses to DNA damage. Genes Dev 16: 560–570.

Koch, B., S. Kueng, C. Ruckenbauer, K. S. Wendt and J. M. Peters, 2008 The Suv39h-HP1 histone methylation pathway is dispensable for enrichment and protection of cohesin at centromeres in mammalian cells. Chromosoma 117: 199–210.

Losada, A., 2008 The regulation of sister chromatid cohesion. Biochim Biophys Acta 1786: 41–48.

Losada, A., M. Hirano and T. Hirano, 1998 Identification of Xenopus SMC protein complexes required for sister chromatid cohesion. Genes Dev 12: 1986–1997.

Losada, A., M. Hirano and T. Hirano, 2002 Cohesin release is required for sister chromatid resolution, but not for condensin-mediated compaction, at the onset of mitosis. Genes Dev 16: 3004–3016.

Mc Intyre, J., E. G. Muller, S. Weitzer, B. E. Snydsman, T. N. Davis et al., 2007 In vivo analysis of cohesin architecture using FRET in the budding yeast Saccharomyces cerevisiae. Embo j 26: 3783–3793.

Merkenschlager, M., and E. P. Nora, 2016 CTCF and Cohesin in Genome Folding and Transcriptional Gene Regulation. Annu Rev Genomics Hum Genet 17: 17–43.

Michaelis, C., R. Ciosk and K. Nasmyth, 1997 Cohesins: chromosomal proteins that prevent premature separation of sister chromatids. Cell 91: 35–45.

Moreno, S., A. Klar and P. Nurse, 1991 Molecular genetic analysis of fission yeast Schizosaccharomyces pombe. Methods in enzymology 194: 795–823.

Nagai, T., A. Sawano, E. S. Park and A. Miyawaki, 2001 Circularly permuted green fluorescent proteins engineered to sense Ca2+. Proc Natl Acad Sci U S A 98: 3197–3202.

Nasmyth, K., 2011 Cohesin: a catenase with separate entry and exit gates? Nat Cell Biol 13: 1170–1177.

Nasmyth, K., and C. H. Haering, 2005 The structure and function of SMC and kleisin complexes. Annu Rev Biochem 74: 595–648.

Osadska, M., T. Selicky, M. Kretova, J. Jurcik, B. Sivakova et al., 2022 The Interplay of Cohesin and RNA Processing Factors: The Impact of Their Alterations on Genome Stability. Int J Mol Sci 23.

Parelho, V., S. Hadjur, M. Spivakov, M. Leleu, S. Sauer et al., 2008 Cohesins functionally associate with CTCF on mammalian chromosome arms. Cell 132: 422–433.

Peters, J. M., A. Tedeschi and J. Schmitz, 2008 The cohesin complex and its roles in chromosome biology. Genes Dev 22: 3089–3114.

Rao, H., F. Uhlmann, K. Nasmyth and A. Varshavsky, 2001 Degradation of a cohesin subunit by the N-end rule pathway is essential for chromosome stability. Nature 410: 955–959.

Remeseiro, S., A. Cuadrado and A. Losada, 2013 Cohesin in development and disease. Development 140: 3715–3718.

Russell, P., and P. Nurse, 1986 cdc25+ functions as an inducer in the mitotic control of fission yeast. Cell 45: 145–153.

Samejima, I., and M. Yanagida, 1994 Bypassing anaphase by fission yeast cut9 mutation: requirement of cut9+ to initiate anaphase. J Cell Biol 127: 1655–1670.

Schär, P., M. Fäsi and R. Jessberger, 2004 SMC1 coordinates DNA double-strand break repair pathways. Nucleic Acids Res 32: 3921–3929.

Schmidt, C. K., N. Brookes and F. Uhlmann, 2009 Conserved features of cohesin binding along fission yeast chromosomes. Genome Biol 10: R52.

Schmiesing, J. A., A. R. Ball, Jr., H. C. Gregson, J. M. Alderton, S. Zhou et al., 1998 Identification of two distinct human SMC protein complexes involved in mitotic chromosome dynamics. Proc Natl Acad Sci U S A 95: 12906–12911.

Schneider, C. A., W. S. Rasband and K. W. Eliceiri, 2012 NIH Image to ImageJ: 25 years of image analysis. Nat Methods 9: 671–675.

Sole-Soler, R., and J. Torres-Rosell, 2020 Smc5/6, an atypical SMC complex with two RING-type subunits. Biochem Soc Trans 48: 2159–2171.

Tanaka, T., M. P. Cosma, K. Wirth and K. Nasmyth, 1999 Identification of cohesin association sites at centromeres and along chromosome arms. Cell 98: 847–858.

Tomonaga, T., K. Nagao, Y. Kawasaki, K. Furuya, A. Murakami et al., 2000 Characterization of fission yeast cohesin: essential anaphase proteolysis of Rad21 phosphorylated in the S phase. Genes Dev 14: 2757–2770.

Tóth, A., R. Ciosk, F. Uhlmann, M. Galova, A. Schleiffer et al., 1999 Yeast cohesin complex requires a conserved protein, Eco1p(Ctf7), to establish cohesion between sister chromatids during DNA replication. Genes Dev 13: 320–333.

Uhlmann, F., F. Lottspeich and K. Nasmyth, 1999 Sister-chromatid separation at anaphase onset is promoted by cleavage of the cohesin subunit Scc1. Nature 400: 37–42.

Uhlmann, F., and K. Nasmyth, 1998 Cohesion between sister chromatids must be established during DNA replication. Current Biology 8: 1095–1102.

Uhlmann, F., D. Wernic, M. A. Poupart, E. V. Koonin and K. Nasmyth, 2000 Cleavage of cohesin by the CD clan protease separin triggers anaphase in yeast. Cell 103: 375–386.

Varshavsky, A., 1996 The N-end rule: functions, mysteries, uses. Proc Natl Acad Sci U S A 93: 12142–12149.

Waizenegger, I. C., S. Hauf, A. Meinke and J. M. Peters, 2000 Two distinct pathways remove mammalian cohesin from chromosome arms in prophase and from centromeres in anaphase. Cell 103: 399–410.

Weitzer, S., C. Lehane and F. Uhlmann, 2003 A model for ATP hydrolysis-dependent binding of cohesin to DNA. Curr Biol 13: 1930–1940.

Xiang, S., and D. Koshland, 2021 Cohesin architecture and clustering in vivo. Elife 10.

Yatskevich, S., J. Rhodes and K. Nasmyth, 2019 Organization of Chromosomal DNA by SMC Complexes. Annu Rev Genet 53: 445–482.

