## Supplementary material for "BiFCo: Visualising cohesin assembly/disassembly cycle in living cells": Supp Figs

### Supplementary Figure1

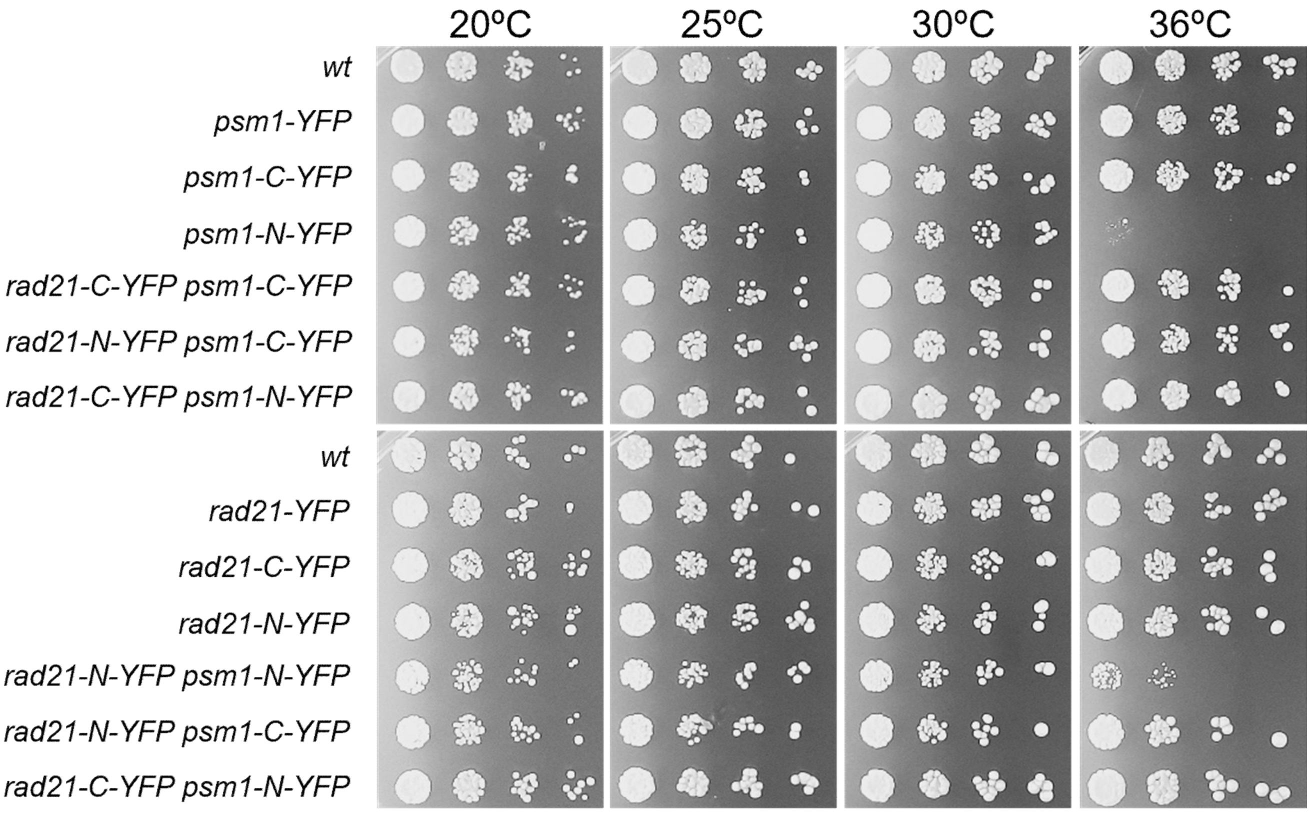


**Supplementary figure 1**. Viability assay. All strains, either single or double-tagged along with a wild type untagged control, where grown to log phase. Number of cells/mL were assessed in a neubauer´s chamber to standardize the number of cells of each strain. Five-fold dilutions were spotted at indicated temperatures to assess viability. Only the fusion of Psm1 with the N-terminal moiety of YFP seems to affect cell viability, albeit only at 36°. It is possible that the spatial conformation that adopts the N-YFP at the C-terminal end of Psm1 hinders the interaction with Psm3 and/or Rad21, only at high temperature, affecting the formation of a functional cohesin ring. In this scenario, reconstitution of the native YFP structure by bi-molecular complementation restores wild-type-like cohesin function. This would also explain why the same N-terminal part of YFP fused to Rad21 does not restore viability

### Supplementary figure 2

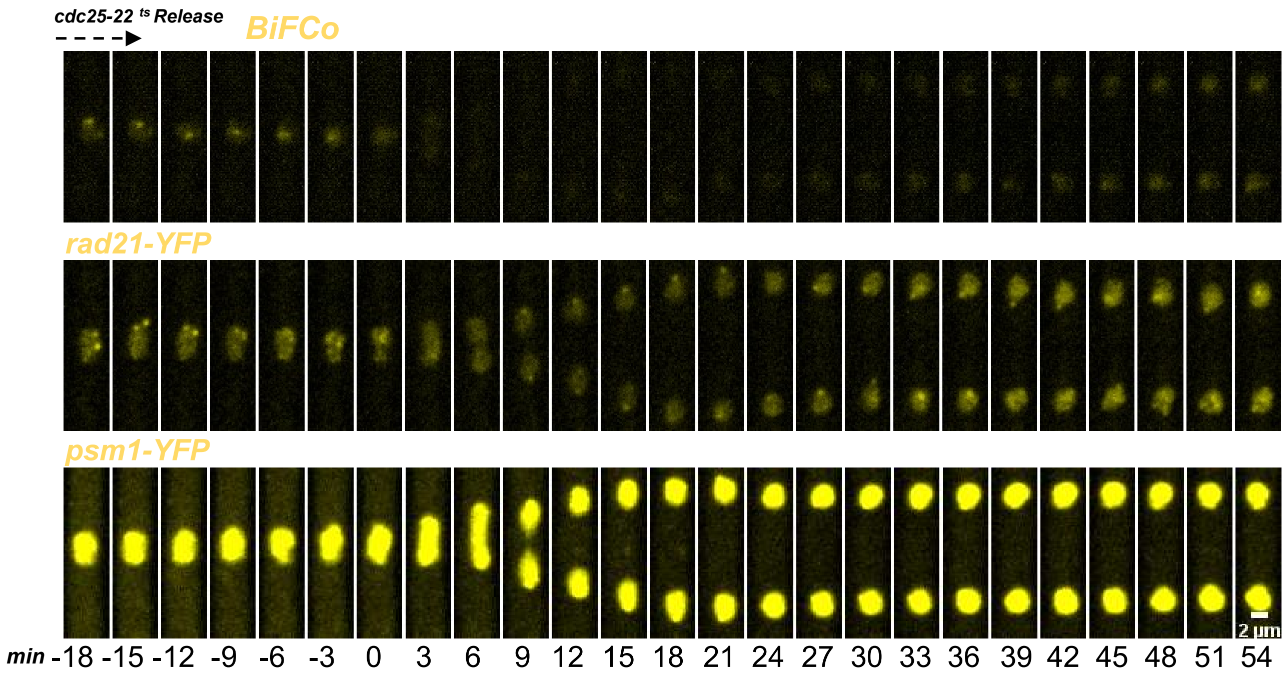


**Supplementary figure 2**. Same frames as in Figure 4A but keeping identical brightness and contrast settings.

**Supplementary Video 1.** BiFCo tracking over two mitotic cycles from figure 3D. cdc25-22 cells were arrested in G2 and released at permissive temperature under the microscope to follow two consecutive mitotic divisions over a total time of 8 hours. Maximal projections every time point (6 minutes) are animated at 4 frames/sec.

**Supplementary Video 2.** Total vs Assembled cohesin components. Time lapse video (3 frames/sec) from the experiment described in figure 4A

**Supplementary Video 3**. BiFCo signal decay in mitosis is dependent on APC/C function. Time lapse video (4 frames/sec) from experiment described in figure 5A
